# Mechanically-gated currents in mouse sensory neurons lacking PIEZO2

**DOI:** 10.1101/2025.08.03.668324

**Authors:** Oscar Sánchez-Carranza, Valérie Bégay, Sampurna Chakrabarti, Mireia Pampols-Perez, Lin Wang, Jonathan Alexis García-Contreras, Annette Hammes, Gary R. Lewin

## Abstract

Touch sensation starts with the opening of mechanically-gated activated ion channels at neuroglial endings of mechanoreceptors in the skin. The function of around half of low threshold mechanoreceptors is dependent on the presence of the mechanically gated ion channel PIEZO2. It has been reported that particularly rapidly-adapting mechanosensitive currents (RA-currents) in the cell bodies of acutely cultured sensory neurons are dependent on PIEZO2. Here we re-examined this question by making a quantitative study of mechanically-gated currents activated by substrate deflection in sensory neurons lacking PIEZO2. We characterized mechanically-gated currents from embryonic and post-natal sensory neurons, taken from global *Piezo2*^*-/-*^ or *Piezo2* conditional knockouts (*Piezo2*^*cko*^), respectively. Surprisingly in both models, *Piezo2* gene deletion was not associated with any significant reduction in the sensitivity or incidence of mechanosensitive currents compared to wild type controls. There was, however, a moderate reduction in the incidence of RA-currents with very fast activation and inactivation kinetics in both embryonic *Piezo2*^*-/-*^ and juvenile *Piezo2*^*cKO*^ mice. These results show that PIEZO2 channels are not the only mechanosensitive channels mediating RA-currents in sensory neurons. Furthermore, our data suggest that the phenotypes associated with *Piezo2* loss of function alleles may sometimes be due to secondary effects of gene deletion, for example, by changing the developmental trajectory of sensory neurons. Emphasis should be put on the diversity of mechanosensitive ion channel function in sensory neurons which needs to be further elucidated.

**Significance:** The mechanosensitive ion channel PIEZO2 has been proposed to be the major mechanically-gated ion channel for the transduction of touch. We used patch-clamp electrophysiology to measure mechanically gated currents in sensory neurons lacking PIEZO2 channels using two distinct genetic strategies. *Piezo2* gene deletion was associated with a moderate reduction in the frequency of very rapidly inactivating mechanosensitive currents in early post-natal sensory neurons. No significant reduction in the sensitivity or incidence of mechanosensitive currents was observed. These data challenge the idea that PIEZO2 channels are the main transducers of force in sensory neurons, at least via substrate deflection. Other interpretations of loss of function phenotypes, including the possibility that early loss of PIEZO2 could alter developmental trajectories of sensory neurons to impair mechanoreceptor function, should be considered.

## Introduction

The mechanically-gated ion channel PIEZO2 has been linked to the senses of touch, proprioception and pain in mammals (1–8). Touch sensation starts with the transduction of mechanical stimuli at the peripheral endings of mechanoreceptors in the skin. It has been shown that for many, but not all mechanoreceptors, the presence of PIEZO2 at sensory endings is absolutely necessary for transduction (1, 2, 9). However, direct measurement of mechanically-gated ion channel activity at sensory endings has so far not been feasible in mammalian preparations (10), but cultured dorsal root ganglion neurons possess mechanically-gated currents postulated to be functionally equivalent to those that mediate transduction at sensory endings (11–15). Indeed, there are several examples where loss of mechanically-gated currents after genetic ablation of transduction candidates is associated with of loss of mechanoreceptor function (1, 2, 15–17). Heterologous expression of PIEZO2 channels in various cells types is associated with mechanically-gated currents with very fast activation and inactivation time constants which have been termed rapidly adapting or RA-currents (13, 14, 18–20). Interestingly, RA-currents measured in cells expressing PIEZO2 have kinetics which are very similar to endogenous mechanosensitive currents in isolated sensory neurons (7, 13, 14). However, even in the case of *Piezo2* genetic deletion in mice where the loss of RA-mechanosensitive currents was described to be particularly profound, many cells still displayed mechanosensitive currents with fast kinetics (1, 2).

Several years ago we developed a technique where mechanosensitive currents can be evoked by moving the cell substrate with single pillar deflections (20, 21). This method allows the quantification of current amplitude in relation to deflection amplitude and reveals mechanically-gated currents with kinetics and biophysical properties almost identical to those produced by cell indentation (7, 17, 20, 22, 23). The mechanically-gated ion channel ELKIN1/TMEM87a was recently identified as being necessary for normal touch receptor function in mice (17, 24). Furthermore, genetic deletion of the *Elkin1* gene was associated with a loss of mechanosensitive currents in cultured sensory neurons evoked with indentation stimuli as well as pili deflection (17). Due to the post-natal lethality of mouse *Piezo2* gene ablation (25, 26) there have been only limited studies on the mechanosensitive currents remaining in sensory neurons lacking PIEZO2 (1, 2, 27). In this study we set out to measure mechanosensitive currents that remain in sensory neurons that genetically lack *Piezo2* in two different mouse models. Our results revealed that complete ablation of the *Piezo2* gene led to very moderate reductions in the incidence of mechanosensitive RA-currents in sensory neurons. Our data strongly suggest that genetic ablation of *Piezo2* can be substantially compensated for by the presence of other mechanically-gated channels in sensory neurons.

## MATERIALS AND METHODS

### Primary cell culture

DRG neurons were collected from all spinal segments (or only lumbar if indicated) in plating medium on ice (DMEM-F12 (Invitrogen) supplemented with L-Glutamine (2 µM, Sigma-Aldrich), Glucose (8 mg/ml, Sigma Aldrich), Penicillin (200 U/mL)-Streptomycin (200µg/mL) and 10 % fetal horse serum). DRGs were treated with Collagenase IV (1 mg/ml, Sigma-Aldrich) for 15 min for embryonic cells (E18.5) and cells from pups (P6), at 37°C and then washed three times with Ca^2+^- and Mg^2+^-free PBS. Samples were then incubated with 0.05% trypsin (Invitrogen, Karlsruhe) at 37°C for 15min. Collected tissue was triturated with a pipette tip and plated in a droplet of plating medium on the silanized elastomeric pillar arrays pre-coated with laminin (4 µg/cm^2^, Invitrogen) as described (*see elastomeric pillar arrays section*) (20, 21). Cells were cultured overnight, and electrophysiology experiments were preformed 18-24 h post-dissection.

### Mice

*Piezo2*^*+/-*^ mice (1, 26) were mated to generate *Piezo2*^*-/-*^ mice (*Piezo2*^*KO*^). The females were monitored for 7 days to observe the presence or absence of vaginal plug during mating. The day when the vaginal plug was observed was considered as the embryonic day 0.5 (E0.5). Females were sacrificed at E18.5 in a 100% CO_2_ chamber. The embryos were collected and put in PBS on ice before DRG extraction as previously described (28). A piece of the tail was cut for genotyping.

*Hoxb8*^*+/Cre*^ (29) mice (Tg(Hoxb8-cre)^1403Uze^; MGI 4881836) were mated with *Piezo2*^*+/-*^ to generate *Hoxb8*^*+/Cre*^;*Piezo2*^*+/-*^ animals. In parallel *Piezo2*^*fl/fl*^ mice (25) were mated with *Ai14*^*f/f*^ mice (30) (Gt(ROSA)^26Sortm14(CAG-tdTomato)Hze^; MGI 3809524) to generate *Piezo2*^*fl/fl*^; *Ai14*^*fl/fl*^. Subsequently, *Hoxb8*^*+/Cre*^; *Piezo2*^*+/-*^ were mated with *Piezo2*^*fl/fl*^; *Ai14*^*fl/fl*^ animals to generate *Piezo2*^*CKO*^ (*Hoxb8*^*+/Cre*^; *Piezo2*^*-/fl*^; *Ai14*^*+/f*^). *Hoxb8*^*+/+*^; *Piezo2*^*-/f*^; *Ai14*^*+/f*^ were used as control animals (*Piezo2*^*Ctrl*^). Littermates were used in each experiment. All experiments with mice were done in accordance with protocols reviewed and approved by the German Federal authorities (State of Berlin).

### Genotyping

To genotype *Piezo2*^*K*O^ and *Piezo2*^*CKO*^ animals, a piece of tail was taken from E18.5 embryos and P1 tattooed pups, respectively. Tissues were incubated overnight at 55° C while shaking at 800 rpm in a lysis buffer containing (in mM): 200 NaCl, 100 Tris pH 8.5, 5 EDTA, 0.2% of SDS and Proteinase K (10 mg/mL, Carl Roth). PCRs were performed using supernatant of the lysis preparation as DNA template (20-100 ng), 1X Taq PRC buffer, 2 mM MgCl_2_, 0.2 mM dNTPs, 1.25U Taq-polymerase (Thermo-fisher Scientific) and 5 pmol of forward- and reverse-specific primers targeting *piezo2* (**Table 1**).

**Table 1.**
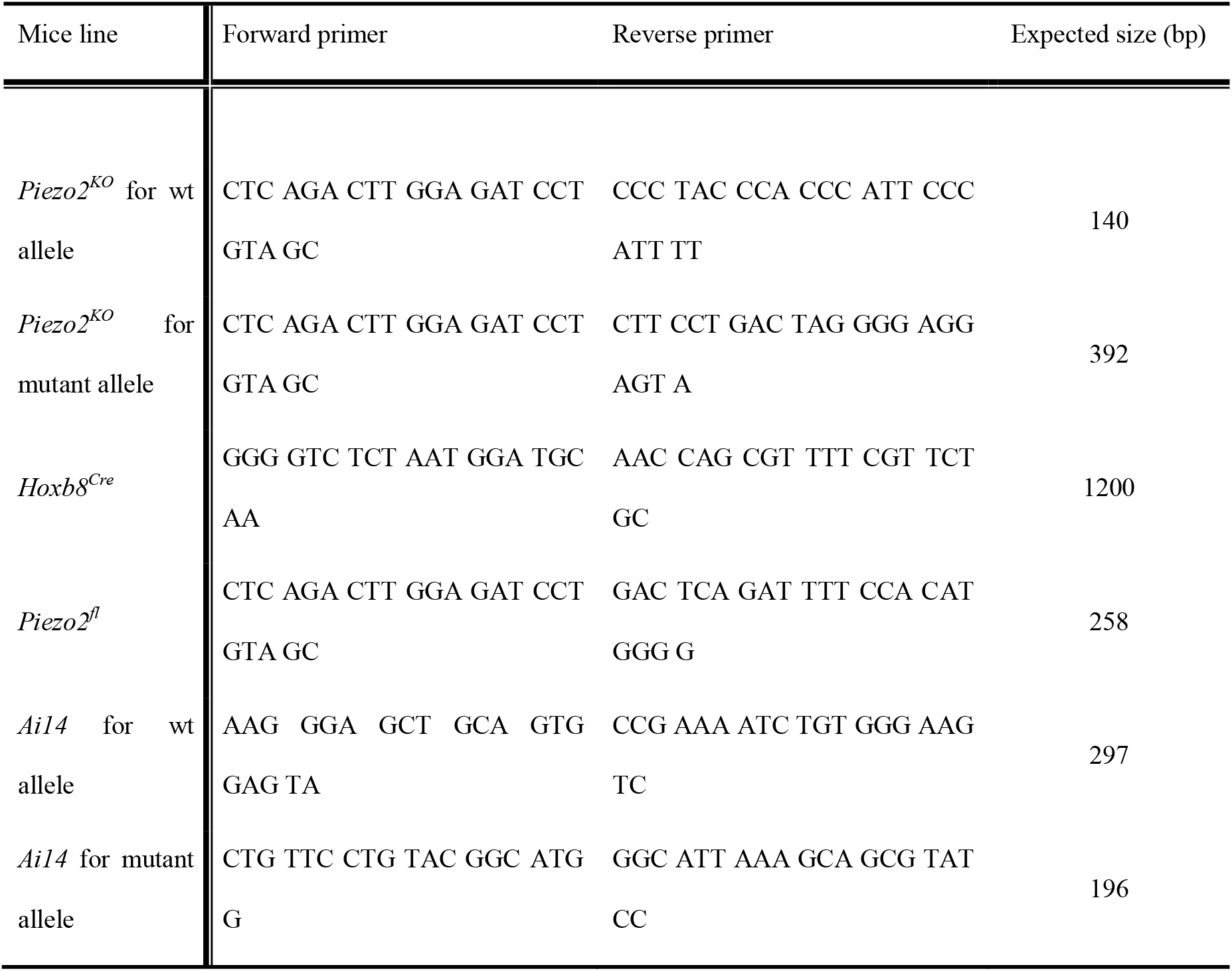
PCR primers for animal genotyping.

### Elastomeric pillar arrays

Elastomeric pillar arrays were prepared as previously described (20). Briefly, freshly mixed PDMS (Sylgard 184, Dow Corning, USA) was prepared in a 1:10 mix of both components and incubated for 30 min in vacuum. Negative masters were covered with PDMS and left for 30 min. Borosilicate glass coverslips (22 × 22 mm, VWR® International, Thickness No.2) were incubated for 2 min in a FEMTO plasma cleaner (Diener Electronic, Nagold, Germany) for oxygen plasma treatment activation and the activated side (upper part) was placed (side down) on the PDMS-covered negative masters. Pillar arrays were incubated at 100° C for 1h. Pillar arrays were pealed apart from the negative masters and were activated by oxygen plasma treatment for 2 min. Each pilus within the array exhibited a radius of 1.79 µm and a length of 5.8 µm. Pillar arrays were silanized using vapour phase (tridecafluoro-1,1,2,2-tetrahydrooctyl) trichlorosilane 97% (AB111444, ABCR GmbH & Co. KG, Karlsruhe, Germany) for 45□min and then coated with laminin (4 µg/cm^2^, Invitrogen) overnight at 37° C in a humid chamber.

### Electrophysiology

Patch clamp experiments were performed at room temperature. Whole-cell patch clamp experiments were carried out in sensory neurons using pulled and heat-polished borosilicate glass pipettes (Harvard apparatus, 1.17 mm x 0.87 mm) with a resistance of 3-6 MΩ. The pipettes were pulled using a DMZ puller (Germany), and filled with a solution containing (in mM): 110 KCl, 10 NaCl, 1 MgCl_2_, 1 EGTA and 10 HEPES. The pH was adjusted to 7.3 with KOH. QX-314 (Alomone Labs) at 1 µM was added to the intracellular solution. The extracellular solution contained (in mM): 140 NaCl, 4 KCl, 2 CaCl_2_, 1 MgCl_2_, 4 Glucose and 10 HEPES. The pH was adjusted to 7.4 with NaOH. Pipette and membrane capacitance were compensated using the auto-function of Patchmaster (HEKA, Elektronik GmbH, Germany) and series resistance was compensated to minimize voltage errors. Currents were evoked by mechanical stimuli (see below) at a holding potential of -60 mV. DRG recordings from *Piezo2*^*+/+*^, *Piezo2*^*+/-*^ and *Piezo2*^*-/-*^ embryos were carried out in a blind manner. When recording neurons from *Piezo2*^*cKO*^, only red cells (td-Tomato+ cells) were selected.

For pillar arrays experiments, a single pilus was deflected using a heat-polished borosilicate glass pipette (mechanical stimulator) driven by a MM3A micromanipulator (Kleindiek Nanotechnik, Germany) as described in (20). Pillar deflection stimuli were applied in the range of 1-1000 nm, larger deflections were discarded. Data were analyzed using Fitmaster software (HEKA Electronik GmbH, Germany). Mechanically-gated currents were classified according to their inactivation kinetics as previously described: the rapidly adapting (RA, τ_inact_ <5 ms), intermediate adapting (IA, τ_inact_ 5-50 ms) and slowly adapting currents (SA, τ_inact_ > 50ms) (20). For quantification and comparison analysis, data was binned by the magnitude of the stimuli (1-10, 11-50, 51-100, 101-250, 251-500, 501-1000 nm) and calculated the mean of the current amplitudes within each bin for every cell. Bright field images (Zeiss 200 inverted microscope) were collected using a 40x objective and a CoolSnapEZ camera (Photometrics, Tucson, AZ) before and during the pillar stimuli to calculate the pillar deflection. The pillar movement was calculated comparing the light intensity of the center of each pilus before and after the stimuli with a 2D-Gaussian fit (Igor Software, WaveMetrics, USA).

### Single molecule Fluorescence in situ hybridization (smFISH)

Lumbar DRGs were collected from P6 pups and were incubated for 40 min in 4% para-formaldehyde washed with PBS and incubated in 30% sucrose (in PBS) overnight at 4°C. DRGs were embedded in OCT Tissue Tek (Sakura, Alphen aan den Rijn). 10 µm-thick cryosections were stored at -80°C until used for experiments. In situ hybridization was carried out according to the manufacturer’s instructions (RNAscopeTM Multiplex Fluorescent V2 assay, ADC, Kit #323110, *piezo2* mouse probe #439971). LSM700 Carls Zeiss and CSU-WI Olympus spinning disk confocal microscopes were used to acquired images at 20X and numerical aperture 0.5 and 0.8, respectively. Fluorescence intensity was analysed using Fiji21.

### Statistical analysis

All data analyses were performed using GraphPad Prism and all data sets were tested for normality. Parametric data sets were compared using a two-tailed, Student’s t-test. Nonparametric data sets were compared using a Mann-Whitney test. To compare more than two groups, One-way ANOVA was used. Categorical data were compared using Fisher’s exact or χ2 tests. Power analyses indicated that the sample sizes were large enough to detect a reduction of 70% in the incidence of MA-currents as previously published (1, 2).

## Results and Discussion

By using cultured sensory neurons from *Piezo2*^*-/-*^ mice we could record mechanically-activated currents (MA-currents) that must have a molecular composition independent of PIEZO2 channels. This experiment was challenging as sensory neurons can only be recorded at the end of embryonic development (E18.5) as *Piezo2*^*-/-*^ mice die shortly after birth (26). We made sensory neuron cultures from all E18.5 embryos obtained from pregnant mice derived from *Piezo2*^*+/-*^ matings and obtained the genotypes using a rapid PCR genotyping protocol (See Methods). Sensory neurons were plated on pillar arrays in order that we could use patch clamp electrophysiology to measure deflection-evoked currents (**Fig. 1A**). The pillar technique is a sensitive way to measure mechanically-gated currents mediated by PIEZO1 and PIEZO2 channels (7, 20, 22, 23) as well as other mechanically gated channels like ELKIN1 and TRPV4 (17, 22, 24). Using indentation stimuli it was shown that the proportion of sensory neurons exhibiting MA-currents increases from around 60% at E13.5 to near 80% at birth (17, 28). To characterize deflection-gated currents in sensory neurons at embryonic stage E18.5, embryonic DRG neurons from the *Piezo2*^*+/*+^, *Piezo2*^*+/-*^ and *Piezo2*^*-/-*^ mice were dissociated and plated on elastomeric pillar arrays pre-coated with laminin. Twenty-four hours after plating, MA-currents were recorded as previously described (7, 17, 20, 22). In cultures from adult DRG we previously showed that around 70% of the cells displayed deflection gated currents (7, 17). In this study we found that only around half of the E18.5 wild type neurons displayed deflection-evoked currents using stimuli between 1-1000 nm, 16/29 neurons from 16 *Piezo2*^*+/+*^ embryos (**Fig. 1B**). Surprisingly, the proportion of responsive cells recorded from *Piezo2*^*-/-*^ embryos (6/12 neurons from 6 embryos) was not different compared to controls or to neurons recorded from *Piezo2*^*+/-*^ embryos (14/24 neurons from 14 embryos). Additionally, the mean amplitudes of deflection-evoked currents for a range of stimuli were also not different between the three genotypes (**Fig. 1C, D**). In responsive cells we observed a reduction in the percentage of deflection stimuli that evoked a current in *Piezo2*^*-/-*^ neurons compared to *Piezo2*^*+/+*^ neurons, yet this small difference was not statistically significant (**Fig. 1E**). As in previous studies using pillar stimuli or indentation (7, 13, 15, 16, 20, 28) sensory neurons displayed deflection gated currents with different inactivation time constants, rapidly-adapting (RA, τ_inact_ <5 ms), intermediately-adapting (IA, τ_inact_ 5-50 ms) or slowly-adapting (SA, τ_inact_ > 50ms) (**Fig. 1F**). For all three genotypes most of the currents recorded were RA, however, these currents made up a substantially smaller proportion of the total in both *Piezo2*^*-/-*^ and *Piezo2*^*+/-*^ sensory neurons which was statistically significant compared to *Piezo2*^*+/+*^ neurons (**Fig. 1F)**. Consistent with this analysis the mean inactivation kinetics (*τ*_inact_) of all mechanically-gated currents recorded from *Piezo2*^*+/-*^ and *Piezo2*^*-/-*^ neurons were 2.5 to 5-fold slower compared to those recorded from *Piezo2*^*+/+*^neurons (**Table2**). The speed of current activation (*τ*_act_) was also significantly slowed in neurons from *Piezo2*^*+/-*^ and *Piezo2*^*-/-*^ embryos as were the latencies for current activation compared to those measured in *Piezo2*^*+/+*^neurons (**Table2**).

**Figure 1.**
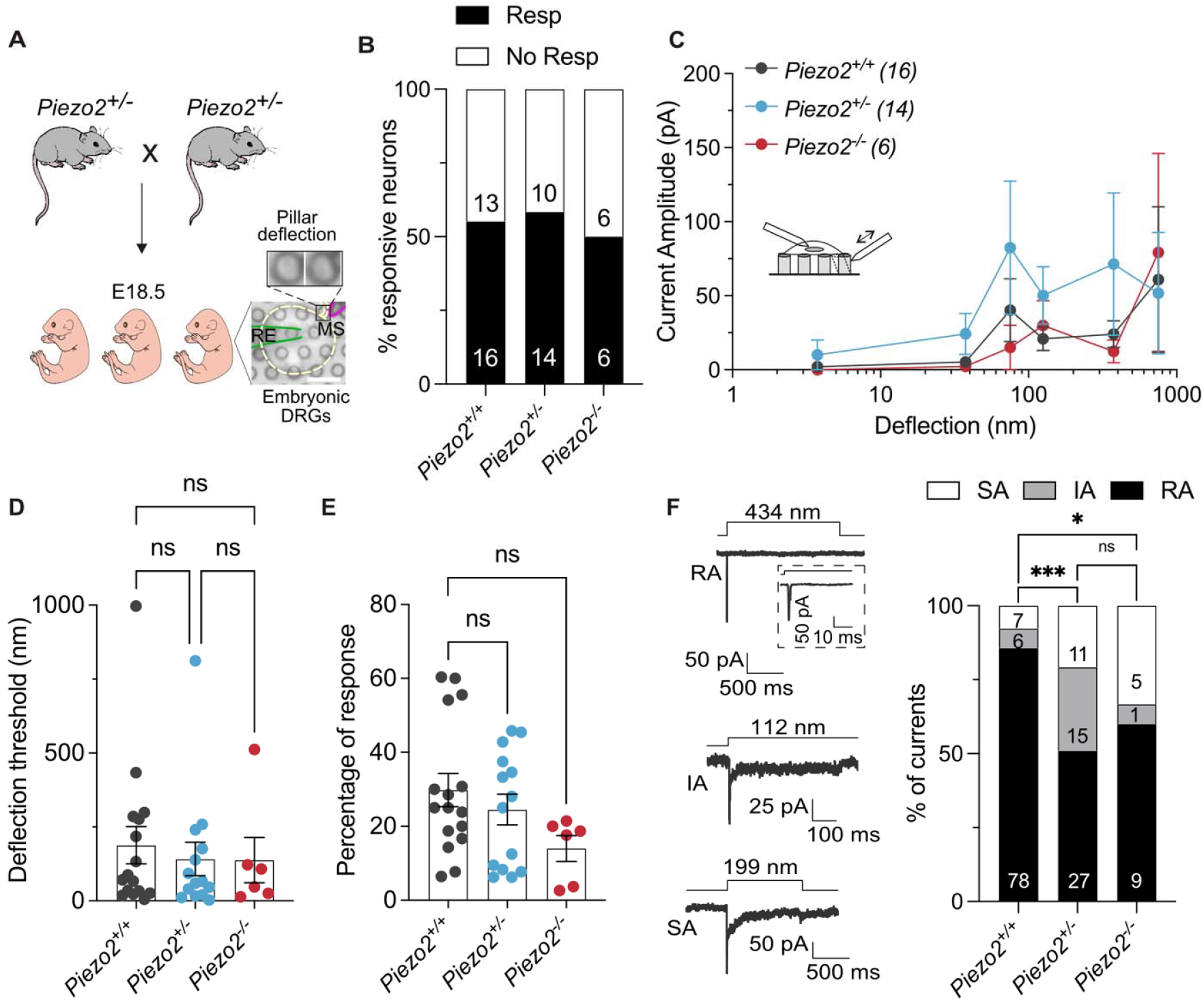
Embryonic DRG neurons from *Piezo2*^*-/-*^ displayed deflection-gated currents. (**A**) Cartoon representing the acquisition of embryonic DRG neurons at E18.5. *Right*, bright field image of an embryonic DRG neuron cultured on pillar arrays. In the insert, the position of a single pilus is shown before and during the deflection stimulus. (**B**) Stacked histogram showing the percentage of responsive (Resp) and non-responsive (No Resp) cells to pillar deflection. Numbers indicate the number of cells. (**C**) Stimulus-response plot of the deflection sensitive currents from embryonic DRGs in *Piezo2*^*+/+*^ (black), *Piezo2*^*+/-*^ (blue) and *Piezo2*^*-/-*^ (red) mice. Data plotted as mean ± s.e.m. (**D**) No differences in deflection thresholds to activate mechanically-gated currents were observed between DRGs from wildtype and mutants. Deflection threshold was calculated as the smallest deflection stimulus applied that evoked a current. Each dot represents one cell. (mean ± s.e.m.) (**E**) The percentage of response was statistically similar in all genotypes. Percentages were calculated according to the total amount of stimulations applied correlate with the stimuli that evoked currents (Kruskal-Wallis test; *Piezo2*^*+/+*^ vs *Piezo2*^*+/-*^, *P*>0.999; *Piezo2*^*+/+*^ vs *Piezo2*^*-/-*^, *P*=0.07). (**F**) *Left*, Representative traces of the three different types of deflection currents in embryonic DRG neurons: RA, IA and SA currents. The insert shows expanded current traces. Deflection stimuli applied are indicated for each trace. *Right*, the percentage of RA currents decreases in DRG neurons from *Piezo2*^*+/-*^ and *Piezo2*^*-/-*^ mice compared to wild type. Values in the histograms indicates the number of the currents observed (Fisher’s exact test; ^***^*P*<0.001, ^*^*P*=0.016).

**Table 2.** Electrophysiological properties of currents recorded from embryonic DRG neurons. *\*P=0.011, **P<0.01, ***P<0.001 (Mann-Whitney test).*

|  | *Piezo2*<sup>+/+</sup> | *Piezo2*<sup>+/-</sup> | *Piezo2*<sup>-/-</sup> |
| --- | --- | --- | --- |
| Cells (no. of currents) | 16 (91) | 14 (53) | 6 (15) |
| Latency (ms) | 2.05 ± 0.19 | 2.32 ± 0.42 | 5.31 ± 0.8*** |
| $t_{\text{act}}$ (ms) | 1.22 ± 0.14 | 2.18 ± 0.4* | 4.16 ± 1.1** |
| $t_{\text{inact}}$ (ms) | 18.65 ± 9.7 | 54.96 ± 15.48*** | 97.4 ± 42.74** |

We were surprised to see no substantial loss of mechanically-gated currents in embryonic sensory neurons as it has been reported that conditional *Piezo2* gene deletion was associated with a substantial loss of RA-currents in adult sensory neurons (1, 2). We thus generated *Piezo2*^*cKO*^ mice by crossing *Hoxb8*^*+/Cre*^;*Piezo2*^*+/-*^ mice with *Piezo2*^*fl/fl*^; *Ai14*^*fl/fl*^ mice to obtain the following genotypes (*Hoxb8*^*+/Cre*^:*Piezo2*^*-/f*^*:Ai14f*^*+/f*^, termed here *Piezo2*^*cKO*^, the control pups had the following genotype (*Hoxb8*^*+/+*^*:Piezo2*^*-/f*^*:Ai14f*^*+/f*.^). In our *Piezo2*^*cKO*^ mice there is an early and complete deletion of the *Piezo2* gene in lumbar sensory neurons which additionally express td-Tomato (**Fig. 2A**). As reported previously, *Piezo2*^*cKO*^ animals started to develop hindlimb movement deficits presumed to derive from loss of proprioception (4). We observed that our *Piezo2*^*cKO*^ animals already at post-natal day 6 (P6) could right themselves from a supine position (**Supp. Video 1**). We verified that in the DRG of *Piezo2*^*cKO*^ virtually no *Piezo2* mRNA was detectable as previously reported (2) (**Fig. 2B**). We made cultures of P6 sensory neurons as our regulatory authorities judged that the burden of the phenotype did not justify the study of these mice at more mature stages. With our strategy *Piezo2*^*cKO*^ sensory neurons showed expression of td-Tomato allowing us to unequivocally identify sensory neurons in culture devoid of PIEZO2. (**Fig. 2A**). Around 60% (19/31 cells from 6 pups) of *Piezo2*^*ctrl*^ neurons showed deflection-evoked currents, but only around 45% (21/46 cells from 6 pups) displayed currents from *Piezo2*^*cKO*^ mice. This small reduction was, however, not statistically significant (Fishers exact test P=0.25) (**Fig. 2C**). As for embryonic sensory neurons, we did not observe differences in the deflection-current amplitude relationship between control neurons and those from *Piezo2*^*cKO*^ mutants (**Fig. 2D,E**).

**Figure 2.**
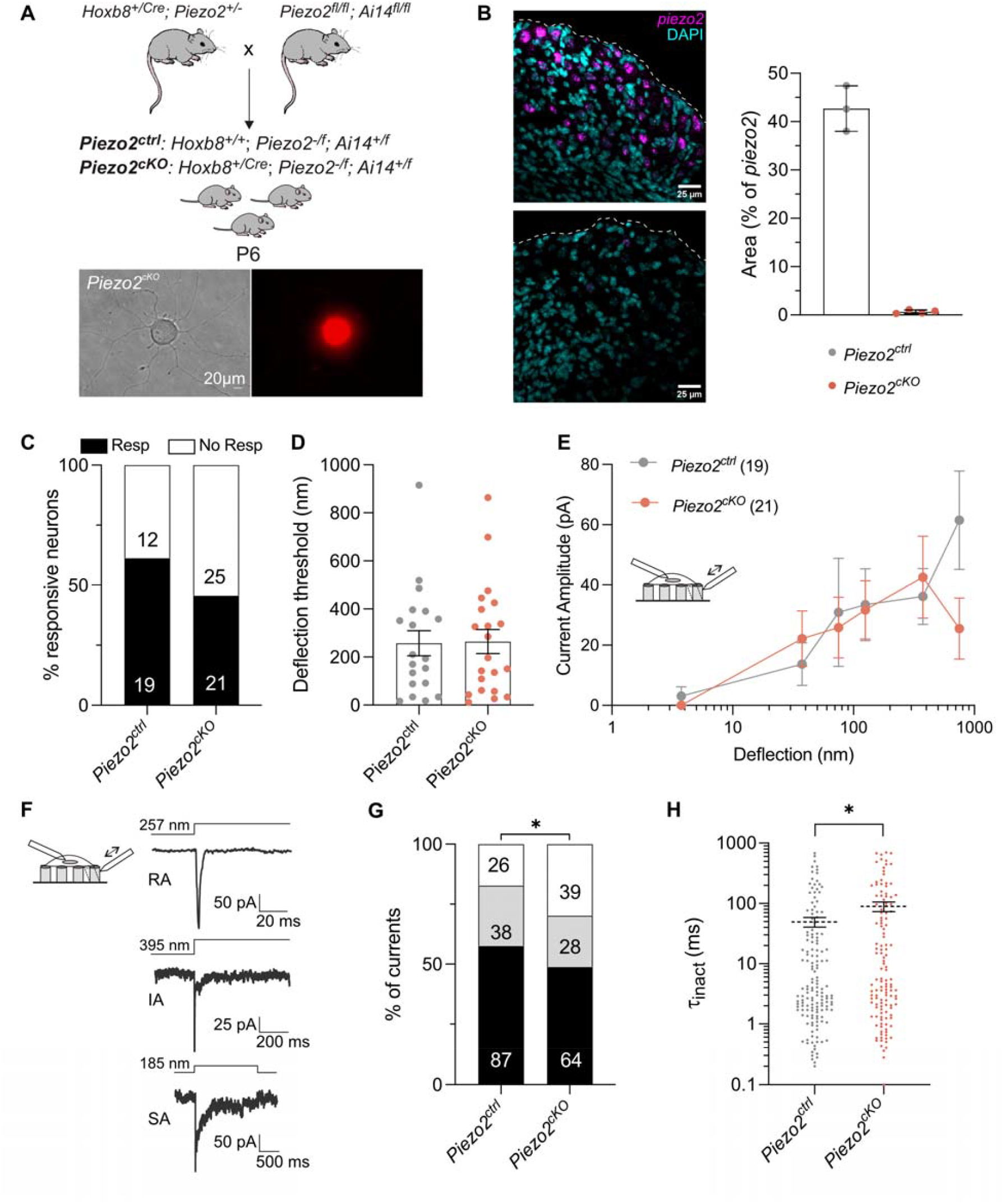
Deflection-gated currents evoked in *Piezo2*^cKO^ neurons. (**A**) *Above*, Scheme for the generation of *Piezo2*^*ctrl*^ and *Piezo2*^*cKO*^ animals. *Below*, acutely prepared dorsal DRG neurons from *Piezo2*^*cKO*^ expressing td-Tomato (red cells). (**B**) *Left*, representative pictures of *piezo2 in situ* hybridization in sections from P6 lumbar dorsal root ganglia. Magenta, *piezo2* mRNA; cyan, 4′,6-diamidino-2-phenylidole (DAPI). Dashed lines show outline of the section Scale bar, 25 µm. *Right*, quantification of area of *piezo2* transcript signal. Each dot represents mean values from several sections from one mouse. (**C**) Stacked histogram showing the percentage of responsive (Resp) and non-responsive (No Resp) dorsal DRG neurons to pillar deflection. Numbers are cells (*P*>0.05, χ^2^ test). (D) The deflection threshold in neurons from *Piezo2*^*cKO*^ mice was similar to controls. (**E**) Stimulus-response plot of the deflection sensitive currents in sensory neurons from *Piezo2*^*ctrl*^ (grey) and *Piezo2*^*cKO*^ mice (salmon). Values are plotted as mean ± s.e.m. (**F**) Representative traces of the three different types of mechanically-gated currents in sensory neurons from P6 animals. RA, IA and SA currents were observed. The deflection stimuli applied are indicated for each trace. The percentage of RA-currents was slightly reduced in neurons from *Piezo2*^*cKO*^ mice. The numbers in the histograms represent the number of the currents observed (*P*=0.04; χ^2^ test). (**G**) Plot showing that τ_inact_ kinetics are slower in *Piezo2*^*cKO*^ compared to controls mouse (Mann-Whitney test, ^*^*P*=0.042).

The three types of currents, RA, IA and SA, were observed in control and mutant cells (**Fig. 2F**). *Piezo2*^*cKO*^ neurons showed a 10% reduction of RA currents compared to control cells (**Fig. 2G**). This again indicates that some RA-currents are *Piezo2*-dependent. Consistent with the reduction in the percentage of RA-currents, we measured more currents in *Piezo2*^*cKO*^ cells with slower inactivation time constants compared to control neurons (**Fig. 2G,H, Table 3**).

**Table 3.**
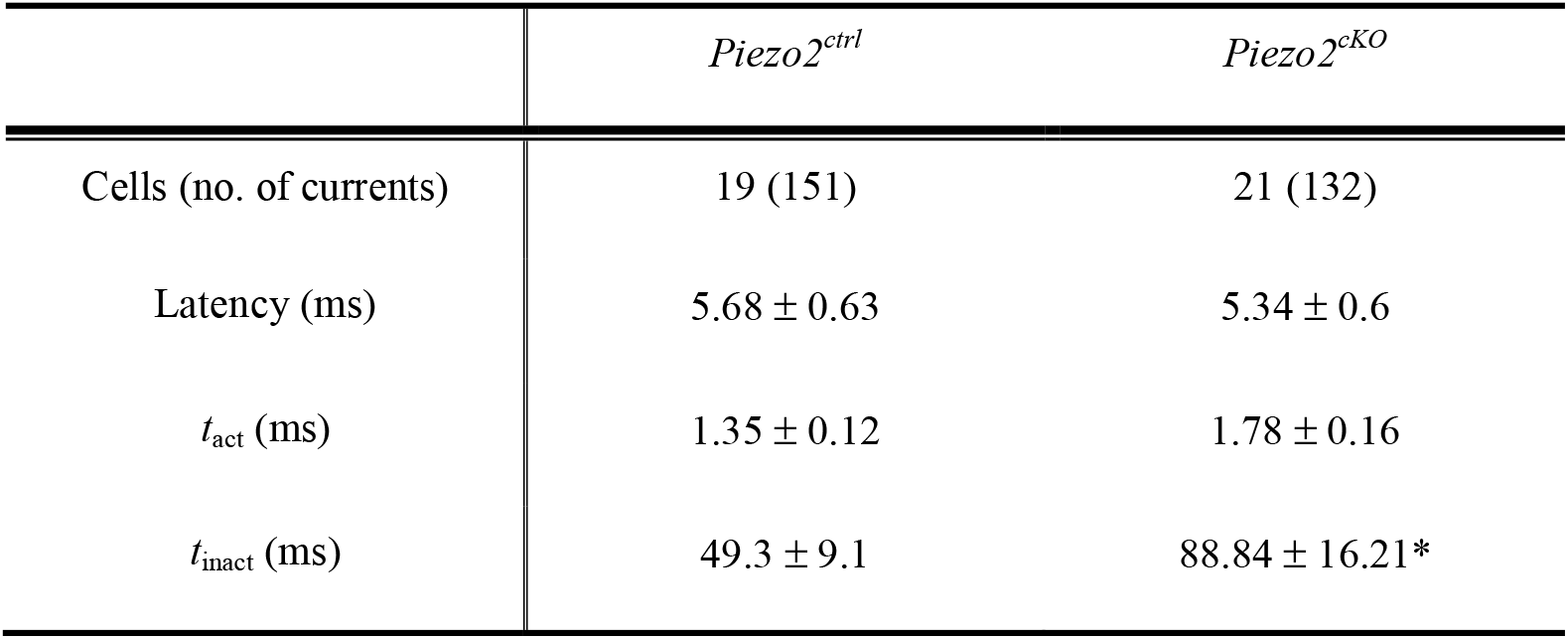
Biophysical properties of currents recorded from *Piezo2*^*cKO*^ DRG neurons. *\*P=0.04 (Mann-Whitney test).*

It has been widely accepted that the PIEZO2 protein is the major transducer of mechanical force in touch receptor neurons (31). Force is transduced at the cutaneous neuroglial endings of mechanoreceptors (32) and genetic ablation of PIEZO2 channels leads to silencing of around 40% of these neurons (1, 2). Loss or silencing of mechanoreceptor function has been associated with loss of mechanosensitive currents in adult cultured sensory neurons (1, 2, 15, 17). Previous studies using two *Piezo2* conditional knockout models also reported a substantial loss of RA-mechanosensitive currents in adult cultured sensory neurons (1, 2). Here we re-examined this issue by recording mechanosensitive currents in late embryonic sensory neurons from *Piezo2* constitutive knockouts as well as in a *Piezo2* conditional knockout model. In contrast to previous reports, we observed no substantial loss of mechanosensitive currents in both models. We did measure a moderate reduction in the incidence of the fastest RA-currents, suggesting that PIEZO2 channels contribute to some of the RA-currents measured in sensory neurons. It is possible that more moderate reductions in the incidence of MA-currents were not detected due to the insufficient sample sizes, although these were similar to previous studies (1, 2). In this study we used the pillar method to measure substrate deflection induced currents, instead of the indentation technique used in earlier studies. However, both indentation and pillar deflection are efficient ways to activate heterologously expressed PIEZO channels (19, 20, 22, 22, 24) and similar mechanosensitive currents are found using both methods in sensory neurons (7, 15, 17, 20). Indeed, recent data shows that these methods applied to study the same mutant mice produce results that are broadly in agreement (7, 15, 17, 20). Another difference between this and earlier studies is that we recorded mechanosensitive currents in late embryonic and early post-natal sensory neurons rather than in adults. It is well documented that mechanoreceptor function matures post-natally (33) and this may be influenced by dynamic changes in expression or function of mechanosensitive channels. The relative lack of effect of *Piezo2* deletion on mechanosensitive currents might suggest that PIEZO2 channels only become important in the mature somatosensory system. However, we observed robust expression of *Piezo2* at P6 (**Figure 2**) and these mice already developed motor abnormalities suggesting that proprioceptor function was already impaired.

The genetic loss of an ion channel can lead to changes in the structure and function of neurons not directly related to loss of ion channel activity. For example, loss of function mutations in the sensory voltage-gated sodium channel Na_V_1.7 are associated with morphological changes in the sensory endings of nociceptors in the skin (34). Mutations that change the biophysical function of ion channels have been used to validate whether the channel contributes to endogenous mechanosensitive channel function (35). In this context, gain of function mutations in the PIEZO2 channel introduced into the mouse were recently shown to have minimal effects on mechanoreceptor function (7) and no effect on proprioceptor function (36). In light of these findings, we should ask why do loss of function mutations lead to such profound changes in touch and proprioception in mice and humans (1, 2, 5). Touch is transduced at the neuroglial endings in the skin, but recent data indicates that sensory Schwann cells actively participate in the transduction of light touch (32, 37). Thus, mechanosensitive channels in the sensory neuron membrane may not be the only determinants of mechanoreceptor function in vivo. Finally, recent work has shown that PIEZO proteins bind MyoD (myoblast determination)-family inhibitor proteins (MDFIC and MDFI) which are transcription factor regulators (38, 39). PIEZO2 sequestration of MDFIC proteins could have significant developmental consequences as pathological variants in the *MDFIC* gene cause perinatal death due to aberrant development of the lymphatic system (39). It was also recently shown with gain and loss of function experiments that PIEZO2 plays a role in coronary artery development (23). Thus, loss of PIEZO2 proteins could in principle alter developmental trajectories of different cell types, including perhaps sensory neurons. In conclusion, our study shows that PIEZO2 is just one of an as yet unknown number of channels that contribute to sensory neuron mechanosensitivity.

## Data availability

Data of this study are available from the corresponding author upon reasonable request.

## Author contributions

Conceptualization: OSC, VB and GRL

Mouse model design, validation and regulatory issues: VB and GRL

Electrophysiology: OSC with the help of SC and JAGC

*In-situ* hybridization: MPP, WL and OSC

Funding acquisition: GRL

Supervision: GRL and AH

Writing original draft: OSC and GRL

Writing review and editing OSC and GRL

## Acknowledgments

We thank Franziska Bartelt, Maria Braunschweig and Kathleen Barda for technical assistance.

## Funding

Deutsche Forschungsgemeinschaft (GRL and AH SFB958-B6)

European Research Council grant to (G.R.L, ERC 789128)

Helmholtz Institute for Translational AngioCardio Science (HI-TAC) to OSC, MPP and AH.

Deutsche Forschungsgemeinschaft (DFG GRK 2318 – 318905415-B1) to AH

## Conflict of interest

The authors declare no competing interests

## Notes

### Competing Interest Statement

The authors have declared no competing interest.

